# Butter: High-precision genomic alignment of small RNA-seq data

**DOI:** 10.1101/007427

**Authors:** Michael J. Axtell

## Abstract

Eukaryotes produce large numbers of small non-coding RNAs that act as specificity determinants for various gene-regulatory complexes. These include microRNAs (miRNAs), endogenous short interfering RNAs (siRNAs), and Piwi-associated RNAs (piRNAs). These RNAs can be discovered, annotated, and quantified using small RNA-seq, a variant RNA-seq method based on highly parallel sequencing. Alignment to a reference genome is a critical step in analysis of small RNA-seq data. Because of their small size (20-30 nts depending on the organism and sub-type) and tendency to originate from multi-gene families or repetitive regions, reads that align equally well to more than one genomic location are very common. Typical methods to deal with multi-mapped small RNA-seq reads sacrifice either precision or sensitivity. The tool ‘butter’ balances precision and sensitivity by placing multi-mapped reads using an iterative approach, where the decision between possible locations is dictated by the local densities of more confidently aligned reads. Butter displays superior performance relative to other small RNA-seq aligners. Treatment of multi-mapped small RNA-seq reads has substantial impacts on downstream analyses, including quantification of *MIRNA* paralogs, and discovery of endogenous siRNA loci. Butter is freely available under a GNU general public license.

## INTRODUCTION

Small RNA-seq is a variant RNA-seq method that captures small RNAs, typically in the ∼15-40 nt size range, via directional RNA ligation, cDNA synthesis, and subsequent sequencing. It is a powerful method for the discovery, annotation, and quantification of small RNA molecules. In eukaryotes, microRNAs (miRNAs), endogenous short interfering RNAs (siRNAs), and Piwi-associated RNAs (piRNAs) are currently the most well-understood and biologically relevant small RNAs. piRNAs are mostly restricted to animal germ-line associated tissues and stem cells (1), endogenous siRNAs are most prominent in plants (2), and miRNAs are ubiquitous in most samples from both plants and animals (3). Small RNAs are smaller than the maximum read-lengths achievable by DNA sequencers, so each small RNA-seq read represents a full-length molecule. Thus it is possible to perform simple quantification of sequences, and in some cases limited *de novo* assembly of small RNA precursors (4, 5), without aligning small RNA-seq data to a reference genome. However, the pattern of small RNA alignments to a reference genome is critical for annotation purposes: Such patterns distinguish the various sub-types from each other and from non-specific RNA fragments resulting from sample degradation. For example, curation of the major database of miRNAs now depends strongly on reference-aligned small RNA-seq data (6). Genomic alignment of small RNA-seq data is thus a critical methodology for the study of small RNAs.

Multi-mapped reads, defined as reads with more than one equally likely alignment position, are a general problem in alignment of all types of RNA-seq data, but especially for small RNA-seq data. One simple solution is to ignore multi-mapped reads entirely. This approach is widely used for canonical RNA-seq data (*e.g.* fragments of polyA+ mRNAs). Relatively long read lengths (∼150 nts), paired-end sequencing, and the fact that most polyA+ mRNAs are inherently non-repetitive minimize the number of multi-mapped reads for canonical RNA-seq data. However, for small RNA-seq, none of these caveats apply, and a great deal of data are lost by ignoring multi-mapped reads altogether. Another simple solution is to place multi-mapped reads randomly at one of the possible positions. This behavior is the default for many popular aligners. However, this method has the obvious drawback that the placement of most multi-mapped reads will be wrong. For instance, a read with two possible locations has a 50% chance of being placed incorrectly, one with three possible locations a 67% chance of incorrect placement, and so forth. A third simple solution is to simply report all possible alignment positions. However, this has the effect of causing alignment files to become excessively large. This method also is prone to mis-interpretation in downstream analyses, because the alignments no longer are representative of abundance, and consequently small RNA production from repetitive areas of the genome will be over-estimated.

More sophisticated approaches to the placement of multi-mapped reads have been described for canonical RNA-seq data derived from fragments of polyA+ mRNAs. Mortazavi et al. (7) described a ‘rescue’ strategy for multi-mapped RNA-seq reads where the densities of uniquely-mapped reads were used to generate a probabilistic framework for placing multi-mapped reads. Other broadly similar recursive-rescue strategies incorporating splice-junction spanning reads have also been described (8, 9). Expectation-maximization modelling can also be utilized to provide robust guidance of multi-mapped RNA-seq reads (10). However, such efforts have been largely focused on efforts to estimate isoform abundances and detect splice-junction spanning reads. Both problems are not applicable to small RNA-seq data, and thus the implementations described above are not appropriate for small RNA-seq analysis. In practice, most previous small RNA-seq analyses have not employed any explicit approach to multi-mapped reads beyond one of the simplistic methods (ignoring them, randomly assigning them, or reporting all possible positions).

piRNAs, endogenous siRNAs, and miRNAs are all processed from larger RNA precursors. To varying extents, processing is heterogenous and results in multiple distinct small RNAs that, when properly mapped to the reference genome, result in a cluster of alignments at the locus of origin. It seems likely that these alignment clusters will often contain some uniquely mapping reads, some reads with just a few possible alignment positions, as well as some reads with a higher number of alignment positions. Thus, a rescue strategy conceptually similar to that described by Mortazavi et al. (7) seems especially well-suited to alignment of small RNA-seq data. Here, I describe the tool butter (Bowtie UTilizing iTerative placEment of Repetitive small rnas), which applies an iterative strategy based on uniquely and low-level multi-mapped reads to increase the precision of small RNA-seq genomic alignments.

## MATERIAL AND METHODS

### Overview of butter

Butter is a perl script that depends upon two other freely available tools: samtools (11) and bowtie (12). Figure 1 summarizes the general alignment methodology used by butter. Users input a reference genome and a set of small RNA-seq reads (in either fasta, fastq, or csfasta formats). Optionally, 3′-adapters may be trimmed off at the user's discretion. For the purpose of bowtie alignment, reads are condensed so that sequences present multiple times in the input are queried just once, while tracking the number of times that sequence was sequenced. After bowtie alignment and during creation of the output alignment file, the unique reads are ‘re-inflated’ so that one alignment per original sequence read is present in the output. This process replaces the original read names; original read names can be retained by disabling this read condensation step with the option -- no_condense, at the cost of increased run times.

**Figure 1.**
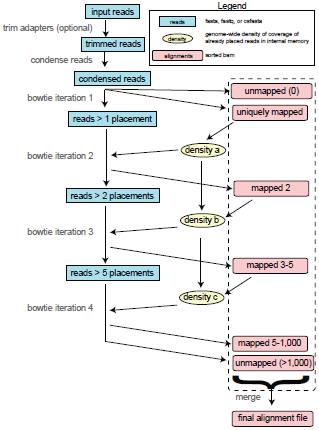
Schematic overview of small RNA-seq alignment by butter.

The first iteration of bowtie limits alignment output to only those reads that have zero or one valid alignment position. Queries with more than one alignment position are written to a temporary fasta or csfasta file for use in the next iteration. The densities of uniquely aligning reads across the entire genome using a sliding window (size=250, step=50) analysis are then calculated and stored to memory. The density in each window is the sum of all of the read-depths at each position (both strands combined) in the window (e.g., ‘area under curve’). The second bowtie iteration uses the multi-mapped reads set aside in the first iteration for a run where only reads with two possible placements are output. Each potential location falls into five different windows. Out of these five, the window with the maximum score is taken as the reference density of each placement. In cases where all possible placements have a reference density of zero, the choice of which placement to retain is simply random. If one possible placement has a non-zero reference density and the other doesn't, the retained placement will be that which is next to the non-zero density. If both possible placements have non-zero reference density, the retained placement is selected based on probabilities dictated by the relative density abundances among the two choices. For example, if placement one had a maximum density score of 70, and placement two had a maximum density score of 30, the probability of placement one being retained is 70 / (70 + 30) [e.g. 70%] and the probability of placement two being retained is 30 / (70 + 30) [e.g. 30%]. As in the first iteration, reads with multi-mappings higher than the set point (two for iteration two) are written to a temporary fasta or csfasta file for analysis in the next iteration. After all reads have been placed, the genome-wide tallies of reference density are updated. After this, the process is repeated for two more iterations. Iteration three captures reads with three, four, or five possible placements, and iteration four captures reads with between five and 1,000 placements (the maximum value of 1,000 can be adjusted with the option --max_rep).

The final output is a single sorted alignment file in the bam format (the binary, compressed representation of the sam format (11)). All adapter-trimmed reads in the input file are represented by a single line in the decompressed alignment, including those that weren't aligned to a genomic position. Besides the custom sam tags added by bowtie, butter adds three custom tags for each alignment. XX:i:[integer] indicates the number of valid alignment positions as reported by bowtie. XY:Z:[string] indicates how the read's final alignment position was chosen by butter, and is one of the following values: U: uniquely mapped, P: multi-mapped and placed due to clustering, R: multi-mapped and randomly placed, N: unmapped, because there were zero valid alignment positions, M: Multi-mappings exceeded setting --max_rep, so no placement performed, O: multi-mapped with no density-based placement possible, number of locations exceeded setting --ranmax, so no placement performed. XZ:Z:[float] indicates the butter-derived probability that was used when placing the read at the reported position. It is set to 1 for reads with an XY:Z: of M, U, N, and O.

### Code availability

The source code for butter is freely available under a GNU general public license at https://github.com/MikeAxtell/butter. Butter is also a required program for ShortStack (3) as of ShortStack version 2.0.0 and is included with the ShortStack package (https://github.com/MikeAxtell/ShortStack).

### Performance analyses using simulated small RNA-seq data

Small RNA-seq data (15 million reads per dataset) were generated using the script sim_srna-seq.pl, version 0.2 (https://github.com/MikeAxtell/sim_srna-seq). Simulated reads were 30% from simulated *MIRNA* loci, 5% from simulated secondary/phased siRNA loci (length 126 nts), and 65% from simulated heterochromatic siRNA loci (length 100 nts). The sim_srna-seq.pl program creates realistic distributions of processing variation from each locus and realistic distributions of abundances. Simulated *MIRNA* and secondary/phased siRNA loci could only derive from non repeat-masked regions of the genome, while simulated heterochromatic siRNA loci could derive from any region of the genome. Supplementary Tables 1 and 2 list complete details on the versions and settings used for all alignments of simulated data, as well as detailed results. Alignments were performed on an iMac running Mac OS 10.9.3 with dual quad-core 3.5GHz processors and 32GB memory (ShortStack 1.2.4, butter, cashx_searchDB, novoalign, bwa, bowtie, and bowtie -m1) or on a linux machine running Ubuntu 12.04.4 LTS with dual quad-core 2.8GHz processors and 32GB memory (patman). Programs that support multi-threading (ShortStack 1.2.4, butter, bwa, bowtie, and bowtie -m1) were run using three processor cores. Strand bias was calculated as the observed number of top-strand matches divided by the expected number of top-strand matches. True positives (TP) were defined as reads placed at the correct position. False positives (FP) were defined as reads placed at the wrong position. False negatives (FN) were defined as reads where a placement was not reported. Sensitivity was defined as TP/(TP+FN), and precision was defined as TP/(TP+FP). The F1 score was calculated as 2TP / (2TP+FP+FN).

### Genome assembly sources

Genome assemblies for *Arabidopsis*, rice, and maize were obtained from Ensembl plants (http://plants.ensembl.org/index.html) on May 23, 2014. Assembly versions were TAIR10 (*Arabidopsis*), IRGSP-1.0 (rice), and AGPv3 (maize).

### Analysis of real small RNA-seq data

Raw fastq files were dumped from sequence read archive accessions SRR1042171.sra; (*Arabidopsis* inflorescence tissue; (13)), SRR976171.sra (rice lamina joints from flag leaves; (14)), and SRR1186264.sra (maize leaves; (15)) using the sra tools function ‘fastq-dump’ under default parameters. Adapter trimming was coincident with alignment by butter by setting the adapter option to CTGTAGGC (SRR1042171 and SRR976171) or ATCTCGTA (SRR1186264). Alignments with butter and bowtie were performed with default options, except that the -m option was set to 1 for the bowtie -m1 alignment. All alignments were processed as needed to produce sorted BAM files. Discovery of clusters dominated by 24 nt siRNAs used a simple script, ‘just_islands.pl’, that defines clusters as regions of continuous small RNA coverage where at least one position exceeds a user-set minimum depth. The minimum depth was set to 20 for these experiments. Clusters where at least 80% of the aligned reads were between 20 and 24 nts in length and for which the most abundant small RNA size was 24 nts were retained. This methodology is similar to that used by ShortStack during it's core *de novo* discovery of small RNA clusters. This script is found as Supplemental Text 1.

## RESULTS

### Butter outperforms all other tested alignment methods in balancing sensitivity with precision

The performance of butter was compared against seven other alignment methods using five sets of simulated small RNA-seq data each from *Arabidopsis*, rice, and maize. Simulated data were used in this analysis because, unlike for real data, the correct alignment location for each simulated read was known with certainty. Thus, the result from each read can be categorized as a true positive (TP; a read aligned to the correct location), a false positive (FP; a read aligned to the incorrect location), or a false negative (FN; a read that was not aligned to the genome). The simulated small RNA-seq data did not include any reads that should not have aligned, so there were no true negatives. The alignment methods (Figure 2A) vary in their treatment of multi-mapped reads. The older version of ShortStack (version 1.2.4; (16)), novoalign, bwa (17), and bowtie (12) all randomly select and report just one alignment position for multi-mapped reads. Patman (18) and cashx_searchDB (19) report all possible alignments for multi-mapped reads. Butter chooses a single alignment position for multi-mapped reads based on the relative densities of unique and confidently placed reads, and also suppresses reporting of any position for multi-mapped reads for which a high-confidence decision can't be made (Figure 1; See methods). Finally, when run with the -m option set to 1, bowtie suppresses all alignments for multi-mapped reads.

**Figure 2.**
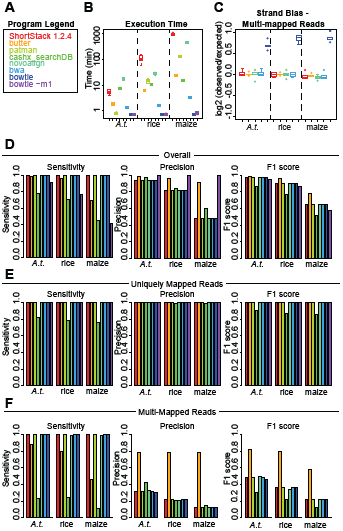
Performance analysis using simulated small RNA-seq data. A. Color-coded legend for the alignment methods used in panels B-F. B. Execution times for alignment of 15 million read simulated small RNA-seq datasets to the indicated plant genomes. Boxplots show the medians, interquartile ranges and outliers for five replicates each. C. Strand bias for placement of multi-mapped reads. Only alignment methods that report a single placement for multi-mapped reads were analyzed. Strand bias is calculated as the log_2_ of the ration between observed top-strand matches and expected top-strand matches. D. Overall sensitivity, precision, and F1 scores. E. As in C except for uniquely mapped reads only. F. As in C except for multi-mapped reads only.

Bowtie, bowtie -m1, and bwa consistently had the most rapid execution times, while ShortStack 1.2.4, patman, novoalign, and cashx_searchDB were typically the slowest (Figure 2B). Butter's execution time was in the middle of pack, and ranked between fourth and fifth out of the tested programs, depending on the species (Figure 2B). Butter's maximum execution time was 24 minutes, for two of the five maize datasets. Among the programs that report just one alignment for each multi-mapped read, bowtie had a very strong bias to reporting alignments to the top genomic strand (Figure 2C), while none of the other programs had a strong bias. Thus, the positions of multi-mapped reads reported by bowtie under default settings are clearly not random, consistent with the warning of strand bias present in the bowtie user manual (http://bowtie-bio.sourceforge.net/manual.shtml). Bowtie with default settings has been a popular method for small RNA-seq alignment. This severe strand bias has thus likely had deleterious impacts on many previous analyses of small RNA-seq data.

As expected, the overall sensitivities (TP/TP+FN) for programs that always report at least one alignment per multi-mapped read were very high, with the exception of cashx_searchDB (Figure 2D). Butter has lower sensitivity, especially for more repetitive genomes (rice and maize), reflecting the fact that it does not report any alignments for the subset of multi-mapped reads where a high-probability position can't be calculated. However, the sensitivity of suppressing reporting of all multi-mapped reads (bowtie -m1) is even lower (Figure 2D). Among methods that attempt to report any alignments for multi-mapped reads, butter displayed superior overall precision (TP/TP+FP; Figure 3C); only by ignoring multi-mapped reads altogether (bowtie -m1) were higher precisions observed. The F1 score (the harmonic mean of sensitivity and precision) was used to assess the overall performance of the alignment methods. Based on this metric, butter had overall performance superior to all other tested methods (Figure 2D). All the methods except cashx_searchDB had virtually identical and nearly perfect performance for uniquely mapped reads (Figure 2E). The differences in performance thus stemmed almost entirely from differences in the placement of multi-mapped reads. For multi-mapped reads, all methods except butter gave very poor precision values, and thus poor F1 scores (Figure 2F). Put in simple terms, most alignments of multi-mapped reads reported by other methods are incorrect, while most alignments of multi-mapped reads reported by butter are correct.

**Figure 3.**
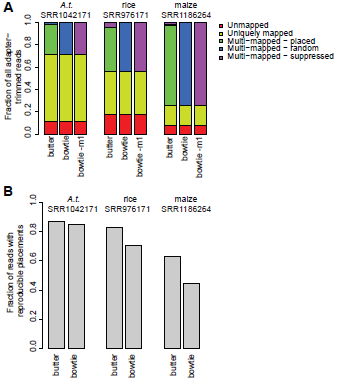
General characteristics of alignments with real small RNA-seq data. A. Overview of alignments of the indicated datasets from *Arabidopsis thaliana* (*A.t.*), rice, and maize with butter, bowtie, and bowtie using option -m set to 1. B. Reproducibility of alignments with bowtie and butter. The indicated datasets were aligned twice with each aligner, and the results to compared to observe the fraction of reads identically placed in both alignments.

### Performance of butter with real data

The impact of alignment method on small RNA-seq data analysis was examined using real data sets from *Arabidopsis* inflorescence tissue (sequence read archive accession SRR1042171; (13)), rice lamina joints from flag leaves (SRR976171; (14)), and maize leaves (SRR1186264; (15)). The depths of these datasets are representative of currently typical small RNA-seq experiments with depths of 14.2, 31.1, and 85.7 million adapter-trimmed reads, respectively. Two commonly used alignment methods were compared with butter: bowtie under default settings, which randomly reports just one position for multi-mapped reads, and bowtie with option -m set to 1, which suppresses all alignments for multi-mapping reads. As expected based on genome structures and sizes, *Arabidopsis* had the fewest number of multi-mapped reads, while maize had the largest (Figure 3A). Strikingly, nearly all multi-mapped reads in all three species could be placed based on clustering density (Figure 3A). It was also apparent that ignoring multi-mapped reads altogether (bowtie -m1) discarded a substantial amount of the data, especially in rice and maize (Figure 3A).

Butter's method of placing multi-mapped reads based on the density of more confidently mapped reads was expected to lessen the variation in alignments due to multi-map position selection, compared to random placement. To test this, the three real data sets were each aligned two times with butter, and two times with bowtie under default settings. The percentage of reads that were identically placed was determined for both butter and bowtie. The more repetitive maize samples generally had the lowest reproducibility, while the least repetitive *Arabidopsis* samples generally had the highest (Figure 3B). In all three species, butter's alignments were more reproducible than bowtie's (Figure 3B).

### Alignment methods can have a large impact on *MIRNA* locus quantification

Identical mature miRNAs are often produced by multiple paralogous *MIRNA* loci. Thus, treatment of multi-mapped small RNA-seq reads is expected to have large effects on the quantification of miRNA production from paralogous loci. As an example, *MIR166* loci that produce identical mature miR166 sequences had radically different numbers of aligned reads depending on the alignment method (Figure 4). In all three species, butter alignments typically indicate one or two dominant paralogs which contribute the majority of miRNA production. In contrast, bowtie alignments result in generally equal accumulation levels across just those loci that happen to be on top genomic strand, and dramatically less for loci that happen to be on the bottom genomic strand (Figure 4). These trends are caused by the random assignment of multi-mapped mature miR166, coupled with the plus strand bias of bowtie's method of multi-mapped position selection (Figure 2C). Finally, suppressing reporting of multi-mapped reads altogether with bowtie -m1 several under-reports accumulation from all loci by omitting the mature miR166 altogether and thus limiting analysis to miRNA-stars and other minor processing variants (Figure 4). The performance analysis (Figure 2) indicated that butter's placement of multi-mapped reads is much more likely to be correct compared to randomly guessing. Thus, quantification of miRNA production from paralogs, and related analyses (such as assessments of *MIRNA* processing precision) are likely to be highly misleading using conventional alignment methods that randomize or ignore multi-mapped reads.

**Figure 4.**
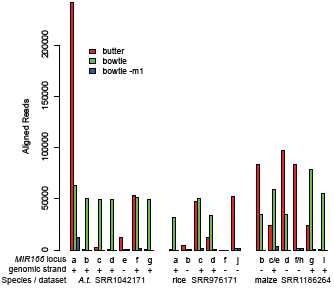
Impact of alignment method on *MIR166* paralog quantification. The indicated *MIR166* paralogs, all of which are annotated as producing an identical miR166 isoform, were quantified from alignments produced by butter, bowtie, and bowtie with the option -m set to 1.

### Alignment methods affect heterochromatic siRNA locus discovery and quantification

Heterochromatic siRNAs are the predominant type of endogenous small RNA in many plants. They are usually 24 nts in length and arise from tens of thousands of genomic loci, many of which are transposons and other types of repeats. Because of their often repetitive nature, treatment of multi-mapped reads during alignment was expected to affect discovery of heterochromatic siRNA-producing loci. Clusters of significant 24 nt siRNA accumulation were determined genome-wide based on the butter, bowtie, and bowtie -m1 alignments of real data. In all three species, the butter alignments shifted the distributions of both cluster sizes and cluster abundances towards larger values, compared with bowtie and bowtie -m1 (Figures 5A–C). Larger cluster sizes and more abundance is an expected consequence of placing multi-mapping reads. For the *Arabidopsis* and rice data, analysis of butter alignments resulted in fewer clusters than bowtie (Figures 5A–B), as expected for an alignment method that tends to build larger clusters. In contrast, for maize, more clusters were found with bowtie alignments than butter (Figure 5C). This is likely due to the greater number of highly multi-mapped maize reads that were suppressed (Figure 3A). Overall, this analysis indicates that alignment methodology indeed does impact the discovery and quantification of heterochromatic siRNAs.

**Figure 5.**
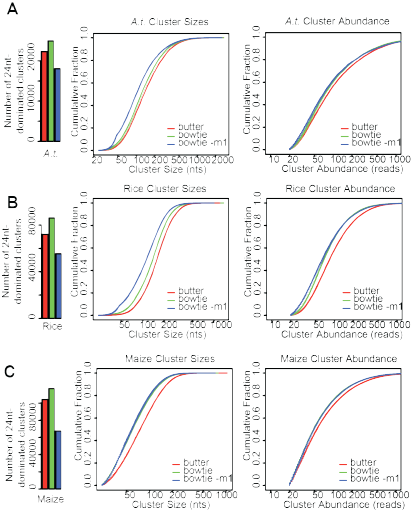
Impact of alignment method on discovery of heterochromatic siRNA loci. A. Left-hand barchart shows the tally of *Arabidopsis thaliana* (*A.t.*) 24 nt siRNA loci discovered from alignment files created by butter (red), bowtie (green), and bowtie with the -m option set 1 (blue). Middle shows cumulative distributions of the sizes of the discovered clusters. Right shows cumulative distributions of the abundances of the discovered clusters. The aligned data was from SRR1042171, as in Figures 3 and 4. B. As in A, except for rice dataset SRR976171. C. As in A, except from maize dataset SRR1186264.

## DISCUSSION

This analysis makes clear that the small RNA-seq alignment methods that have been standard in the field represent rather drastic trade-offs between sensitivity and precision. Randomly choosing a single position for a multi-mapped read preserves high sensitivity at the cost of large reductions in precision, while ignoring multi-mapped reads altogether preserves high precision at the cost of large reductions in sensitivity. These trade-offs become more severe with increasing genome complexity and overall repetitiveness (for instance, the trade-offs are much more severe in maize relative to *Arabidopsis*). Butter is a moderate intermediate between these two extremes, and does a better job at balancing sensitivity (by placing the sub-set of multi-mapped reads where a high-probability location can be found) with precision (by suppressing the reporting of multi-mapped reads where no high-probability location can be found).

Another important observation is that the popular bowtie aligner, when run under the default settings, has a strong strand-bias in its selection of which position to report for multi-mapped reads (Figure 2C). Indeed, this is a known issue with bowtie discussed in its user manual (http://bowtie-bio.sourceforge.net/manual.shtml). Nonetheless, bowtie with default settings has been widely used to align small RNA-seq data. Because strand specificity is a key component of siRNA, miRNA, and piRNA annotations, this systematic error may have had negative consequences on previous work. At a minimum, it is important that users be aware of, and take steps to mitigate this issue when using bowtie under default settings. Note that although butter uses bowtie as its core alignment engine, it's control of bowtie does not allow bowtie to decide on the final placement of multi-mapped reads, and the strand bias for multi-mapped reads is mitigated (Figure 2C).

This analysis also makes clear that the choice of how to deal with multi-mapped small RNA-seq reads has consequences for downstream analysis for both miRNAs and heterochromatic siRNAs. For miRNAs, quantification of production from paralogous genes that produce identical mature miRNAs is strongly dependent on the treatment of multi-mapped reads. In particular, using random placement is highly likely to ‘gloss over’ substantial differences in expression between paralogs. Random placement, or ignoring multi-mapped reads altogether, also tend to make discovered 24 nt heterochromatic siRNA clusters appear smaller and less abundant. Although this analysis focused on plant data, there is no reason to suspect these general observations won't apply for animal and human small RNA-seq data as well, and butter's methods are not specific to any single type of organism.

Overall, this analysis indicates that alignment methods for small RNA-seq can have large impacts on all subsequent analyses. The program butter offers one solution to increase performance in this arena, and it is hoped that this tool assists with future research. More broadly, this analysis highlights the need for more careful consideration of small RNA-seq alignment methods, particularly with respect to multi-mapped reads. Research involving reference-aligned small RNA-seq data should ideally always include a careful description of the alignment methods used including how multi-mapped reads were treated. In addition to the standard deposition of raw data in public archives (GEO, SRA), submission of alignment files themselves should also be encouraged.

## ACKNOWLEDGEMENTS

I thank all members of the Axtell Lab for their comments on the manuscript, and Alice Lunardon, whose data and thoughts provided the initial motivation for me to explore this area of research.

## FUNDING

This work was supported by the National Science Foundation [grant number 1339207 to M.J.A.]. Funding for open access charge: National Science Foundation [grant number 1339207 to M.J.A.].

## REFERENCES

1. Juliano, C., Wang, J. and Lin, H. (2011) Uniting germline and stem cells: the function of Piwi proteins and the piRNA pathway in diverse organisms. Annu. Rev. Genet., 45, 447–469.

2. Coruh, C., Shahid, S. and Axtell, M.J. (2014) Seeing the forest for the trees: annotating small RNA producing genes in plants. Curr. Opin. Plant Biol., 18, 87–95.

3. Axtell, M.J. (2013) ShortStack: comprehensive annotation and quantification of small RNA genes. RNA, 19, 740–751.

4. Wu, Q., Luo, Y., Lu, R., Lau, N., Lai, E.C., Li, W.-X. and Ding, S.-W. (2010) Virus discovery by deep sequencing and assembly of virus-derived small silencing RNAs. Proc. Natl. Acad. Sci. U. S. A., 107, 1606–1611.

5. Kreuze, J.F., Perez, A., Untiveros, M., Quispe, D., Fuentes, S., Barker, I. and Simon, R. (2009) Complete viral genome sequence and discovery of novel viruses by deep sequencing of small RNAs: A generic method for diagnosis, discovery and sequencing of viruses. Virology, 388, 1–7.

6. Kozomara, A. and Griffiths-Jones, S. (2014) miRBase: annotating high confidence microRNAs using deep sequencing data. Nucleic Acids Res., 42, D68–73.

7. Mortazavi, A., Williams, B.A., McCue, K., Schaeffer, L. and Wold, B. (2008) Mapping and quantifying mammalian transcriptomes by RNA-Seq. Nat. Methods, 5, 621–628.

8. Cloonan, N., Xu, Q., Faulkner, G.J., Taylor, D.F., Tang, D.T.P., Kolle, G. and Grimmond, S.M. (2009) RNA-MATE: a recursive mapping strategy for high-throughput RNA-sequencing data. Bioinformatics, 25, 2615–2616.

9. Bonfert, T., Csaba, G., Zimmer, R. and Friedel, C.C. (2012) A context-based approach to identify the most likely mapping for RNA-seq experiments. BMC Bioinformatics, 13, S9.

10. Li, B., Ruotti, V., Stewart, R.M., Thomson, J.A. and Dewey, C.N. (2010) RNA-Seq gene expression estimation with read mapping uncertainty. Bioinformatics, 26, 493–500.

11. Li, H., Handsaker, B., Wysoker, A., Fennell, T., Ruan, J., Homer, N., Marth, G., Abecasis, G., Durbin, R. and 1000 Genome Project Data Processing Subgroup (2009) The Sequence Alignment/Map format and SAMtools. Bioinformatics, 25, 2078–2079.

12. Langmead, B., Trapnell, C., Pop, M. and Salzberg, S.L. (2009) Ultrafast and memory-efficient alignment of short DNA sequences to the human genome. Genome Biol., 10, R25.

13. Creasey, K.M., Zhai, J., Borges, F., Van Ex, F., Regulski, M., Meyers, B.C. and Martienssen, R.A. (2014) miRNAs trigger widespread epigenetically activated siRNAs from transposons in Arabidopsis. Nature, 508, 411–415.

14. Wei, L., Gu, L., Song, X., Cui, X., Lu, Z., Zhou, M., Wang, L., Hu, F., Zhai, J., Meyers, B.C., et al. (2014) Dicer-like 3 produces transposable element-associated 24-nt siRNAs that control agricultural traits in rice. Proc. Natl. Acad. Sci. U. S. A., 111, 3877–3882.

15. Diez, C.M., Meca, E., Tenaillon, M.I. and Gaut, B.S. (2014) Three groups of transposable elements with contrasting copy number dynamics and host responses in the maize (Zea mays ssp. mays) genome. PLoS Genet., 10, e1004298.

16. Shahid, S. and Axtell, M.J. (2014) Identification and annotation of small RNA genes using ShortStack. Methods, 67, 20–27.

17. Li, H. and Durbin, R. (2009) Fast and accurate short read alignment with Burrows-Wheeler transform. Bioinformatics, 25, 1754–1760.

18. Prüfer, K., Stenzel, U., Dannemann, M., Green, R.E., Lachmann, M. and Kelso, J. (2008) PatMaN: rapid alignment of short sequences to large databases. Bioinformatics, 24, 1530–1531.

19. Fahlgren, N., Sullivan, C.M., Kasschau, K.D., Chapman, E.J., Cumbie, J.S., Montgomery, T.A., Gilbert, S.D., Dasenko, M., Backman, T.W.H., Givan, S.A., et al. (2009) Computational and analytical framework for small RNA profiling by high-throughput sequencing. RNA, 15, 992–1002.

